# Chaotic propagation of spatial cytoskeletal instability modulates integrity of podocyte foot processes

**DOI:** 10.1101/065839

**Authors:** Cibele V. Falkenberg, Evren U. Azeloglu, Mark Stothers, Thomas J. Deerinck, Yibang Chen, John C. He, Mark H. Ellisman, James C. Hone, Ravi Iyengar, Leslie M. Loew

**Affiliations:** R. D. Berlin Center for Cell Analysis & Modeling, U. Connecticut School of Medicine, Farmington, CT; Department of Pharmacological Sciences, Icahn School of Medicine at Mount Sinai, New York, NY; Department of Mechanical Engineering, Columbia University, New York, NY; National Center for Microscopy and Imaging Research, UCSD, San Diego, CA; Division of Nephrology, Icahn School of Medicine at Mount Sinai, New York, NY

**Keywords:** podocytes,, actin cytoskeleton,, stability analysis,, serial block-face SEM,, dynamical modeling;, partial differential equations-based modeling.

## Abstract

The kidney podocyte’s function depends on its distinctive morphology. Each podocyte has fingerlike projections, called foot processes, that interdigitate with the processes of neighboring cells to form the glomerular filtration barrier. The integrity of foot process interactions depends on tight spatial control of the dynamics of the underlying actin cytoskeleton, which is regulated by the GTPases, Rac1 and RhoA. To understand how spatially-specific regulation of actin filament dynamics within foot processes controls local morphology, we used a combination of 3-D microscopy and dynamical models. We experimentally determined cell-cell interactions using serial blockface scanning electron microscopy and reconstructed a 3-D spatial representation of a podocyte. We developed a minimal dynamical system for regulation of the actin cytoskeleton; using this 3-D model, we determined how spatial reaction-diffusion dynamics can dysregulate actin bundling, leading to propagation of chaotic foot process effacement. Consistent with experimental observations, our simulations predicted that hyperactive RhoA could destabilize the cytoskeleton. Our simulations showed that deleterious mechanochemical stimuli could lead to local heterogeneity of cytoskeletal dynamics resulting in the emergence of progressive and chaotic loss of foot processes. While global enhancement of Rac1 may result in stronger bundles, the spatial simulations showed that even transient local heterogeneities in polymerization could have dramatic consequences in the stability of multiple foot processes. We conclude that the podocyte morphology optimized for filtration contains intrinsic fragility whereby local imbalances in biochemical and biophysical reactions lead to morphological changes associated with glomerular pathophysiology.

## Introduction

Podocytes, visceral epithelial cells of the kidney glomerulus, enable the selectivity of the glomerular filtration barrier through their specialized morphology. The cytoskeleton of each highly differentiated podocyte is composed of F-actin, microtubules, and intermediate filaments. All three of these cytoskeletal polymers form the cell body and primary processes, but only actin shapes the foot processes (FPs), the delicate fingerlike projections that interdigitate to form the glomerular filtration barrier (1). Actin is organized in a spatially specific fashion. Polymerization beneath the plasma membrane gives rise to a cortical actin network, and addition of crosslinkers result in the high density longitudinally aligned bundles, found in the center of the FPs (2). The FPs establish contact between podocytes and the glomerular basement membrane, in addition to the cell-cell contact via specialized transmembrane junctions called the slit diaphragm, where plasma is filtered (3). As a result of their highly dynamic filtration function, each FP must withstand tensile stresses (due to glomerular expansion during systole) and transverse shear stresses (imposed by the fluid flow crossing the slit diaphragm), while maintaining contact with the neighboring cells as well as the basement membrane (4, 5). The loss of the characteristic FP morphology (i.e., foot process effacement) and reduction in the number of podocytes are common markers of chronic kidney disease (CKD), in which glomerular filtration is compromised (6, 7). The limited treatment options for CKD are in part due to the sensitive nature of these highly differentiated cells.

Podocytes rapidly dedifferentiate after isolation of glomeruli, losing expression of key slit diaphragm proteins and the specialized morphology within 8 hours, and fully reverting to amorphous epithelial cell morphology within 48 hours (8). Cultured primary or immortalized podocytes fail to fully differentiate (9). These observations suggest that the mechanical stresses experienced by the podocyte in the mammalian glomerulus may regulate the integrity of the actin cytoskeleton within the FPs. Without the *in vivo* mechanical stimuli and the final shape signals, cultured podocytes present geometric characteristics (e.g., surface-to-volume ratio, eccentricity, characteristic length, etc.) that are clearly different from the *in vivo* structure (10, 11). Consequently, it is known that protein localization, gene expression levels, and active signaling pathways in cultured podocytes are not similar those of the *in vivo* system. Slit diaphragm has not yet been reconstituted in culture; hence, the unique localization of podocyte specific proteins that lead to the pro-differentiation signaling landscape and the stable actin cytoskeleton of the FPs is absent in cultured podocytes *in vitro* (12, 13). Therefore, in order to understand how the maintenance of the FP morphology and slit diaphragm integrity is regulated, it is important to study the balance of signals within the context of the *in vivo* cell morphology.

Small Rho GTPases play important roles in the regulation of the actin cytoskeleton (14). Rac1 promotes the formation of a branched actin network, as found in lamelipodia, whereas, RhoA promotes the formation of stress fibers and actin bundles. Expression and activity of both of these GTPases are tightly regulated in the healthy podocyte (15). For example, while Rac1 knockdown may prevent protamine sulfate-driven FP effacement (a standard animal model of acute podocyte injury), suggesting increased FP stability; Rac1 knockout animals that are subjected to chronic hypertension exhibit FP loss, albuminuria and glomerulosclerosis, showing an opposite effect (16). Proteinuria and focal FP effacement are also observed when Rac1 is hyperactive (17). It is not clear how an actin polymerization signaling hub, such as Rac1, may act both as a stabilizing and destabilizing factor under these varying mechanical conditions. It is also puzzling that similar phenotypes could be observed for both Rac1 knockdown and overexpression. A similar outcome is observed with RhoA activity. *In vivo* induction of constitutively active or inactive RhoA damages the actin cytoskeleton causing loss of foot processes and albuminuria (18, 19). These finding indicate that the levels and activities of both RhoA and Rac1 need to be within a defined range to maintain foot process integrity. What is the quantitative relationship between local morphology and the GTPase regulators of biochemical and biophysical reactions underlying actin cytoskeleton dynamics that stabilizes podocyte FPs? To answer this question we need to develop computationally tractable models based on realistic *in situ* morphologies. We developed dynamical models that combine upstream GTPase signaling with mechanical forces that control the podocyte actin cytoskeleton. Our model accounts for the actin stoichiometry, exchange between monomeric, fiber, or bundled states, and the *in vivo* podocyte morphology. To account for the spatial specificity of the complex 3-D geometry of the FPs *in vivo,* we constructed a new quantitative model of a representative podocyte using 3-D serial blockface scanning electron microscope (SBEM) imaging in healthy rats. Our 3-D reconstructed model has sufficient resolution to capture the relationship between volume, surface area and characteristic length of the podocyte geometric features as well as the underling biochemical and biophysical reactions. Our cytoskeleton model and the spatially specific simulations provide a mechanistic understanding of the *in vivo* observations regarding Rac1 and RhoA dynamics in podocytes, and their relationship cytoskeletal stability of FPs. The spatial simulations reveal the emergence of chaotic spatial heterogeneities within the actin cytoskeleton when Rac1/RhoA balance is altered providing a multiscale mechanism for foot process effacement. Using this dynamical model, we also show how compensatory mechanisms may impact podocyte cytoskeletal integrity when they appear at different stages of regulatory dynamics.

## Results

### Podocyte morphology

Our first objective was to obtain reconstructed 3-D geometry of interacting podocyte that would sufficiently identify and represent the key spatial features of the podocyte to permit construction of dynamical models. SBEM image stacks processed by manual segmentation and Gaussian filtering resulted in extraordinarily detailed reconstruction of podocytes with arborized processes that included nanoscale cell-cell junctions at the foot processes (Supplementary Movie 1). These reconstructions revealed a complex network of primary branches emanating from the cell body (Supplementary Material 1). Branching angles and the extent of secondary branching for these processes varied; however, the length and number of processes were consistent among different podocytes (Fig S1). Similarly, despite the large level of deviations in the shape and projection pattern for primary processes, other key geometric characteristics, such as volume, principal dimensions, and surface area of individual podocytes exhibited remarkably low variability between individual podocytes (Table 1). Even though SBEM was recently used to reconstruct qualitative features of podocytes (20), to the best of our knowledge, our study represents the first quantitative cataloging and reconstruction of the adult mammalian podocyte geometry *in situ.* The results from the analysis of five different healthy rat podocyte cells are summarized in Table 1, with the steps required to achieve this analysis illustrated in Fig 1..

**Table 1.** Geometric properties, such as surface area (A) and volume (V), of healthy adult rat podocytes are categorized according to different segments of the cell, namely: cell body (CB), major processes (MP), and foot processes (FP) of each rat podocyte (RP). The last column outlines the quantitative morphometric parameters used to generate the idealized, representative podocyte geometry.

| Cell | RP1 | RP8 | RP9 | RP11 | RP13 | Mean | St. D. | Idealized |
| --- | --- | --- | --- | --- | --- | --- | --- | --- |
| $A_{Co+MP} / V_{Co+MP}, \mu^{-1}$ | 1.8 | 1.9 | 1.9 | 1.5 | 1.5 | 1.7 | 0.2 | 1.7 |
| $A / V, \mu m^{-1}$ | 3.2 | 3.2 | 3.4 | 2.6 | 3.0 | 3.1 | 0.3 | 3.2 |
| $A_{FP} / V_{FP}, \mu m^{-1}$ | 8.2 | 8.6 | 8.8 | 8.4 | 9.3 | 8.7 | 0.4 | 9.0 |
| $(A - A_{CB+MP}) / A_{CB+MP}, \%$ | 130 | 110 | 130 | 100 | 150 | 120 | 20 | 140 |
| $V_{FP} / V, \%$ | 22 | 19 | 21 | 16 | 20 | 20 | 2 | 20 |
| No of branches | 43 | 30 | 34 | 31 | 34 | 34.4 | 5.1 | 36 |
| Length of branches | 10.5 | 10.4 | 12.0 | 11.2 | 10.7 | 11.0 | 0.7 | - |
| $V_{CB} / V, \%$ | 38 | 33 | 21 | 46 | 50 | 38 | 11 | - |
| $V_{MP} / V, \%$ | 40 | 47 | 57 | 38 | 31 | 43 | 10 | - |
| $V_{Nu} / V_{CB}, \%$ | 24 | 42 | 42 | 24 | 22 | 31 | 10 | - |

**Fig 1.**
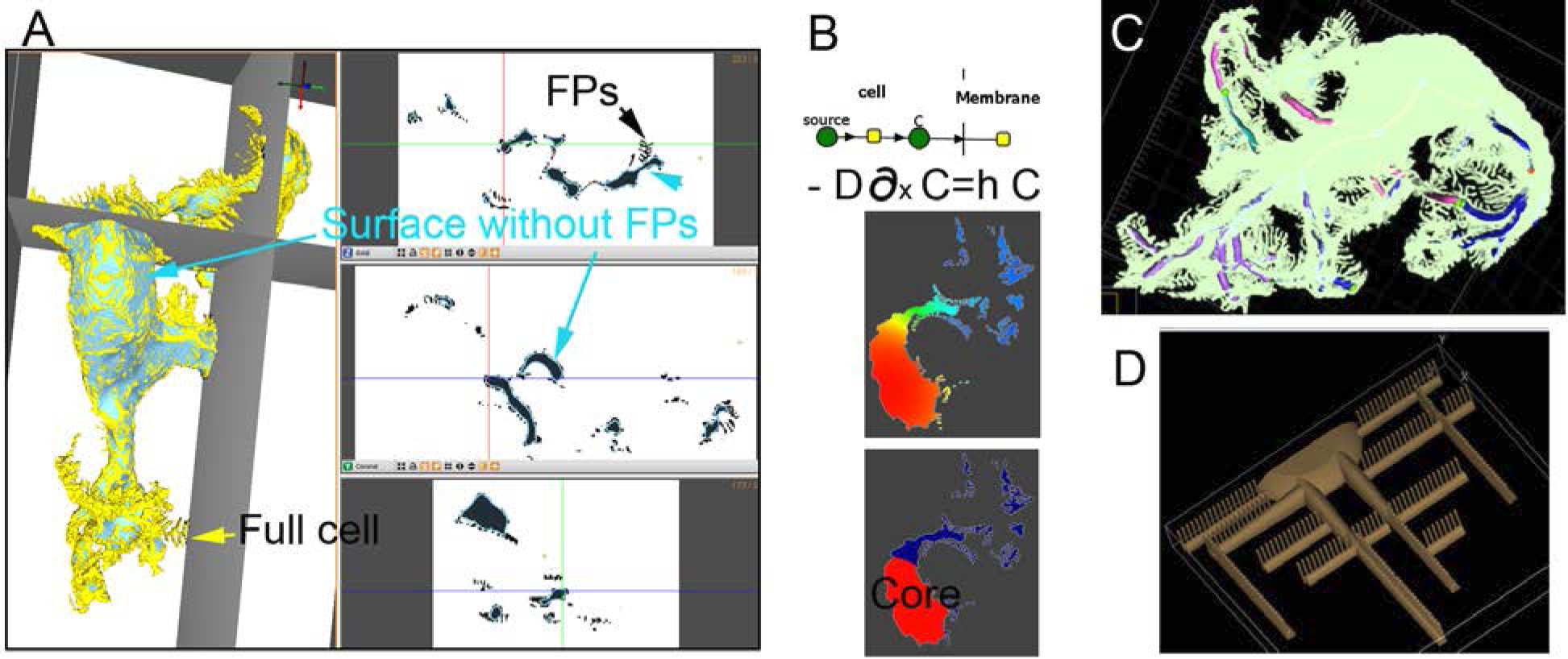
Analysis of podocyte morphology. **(A**) Quantification of contribution of FPs to cell volume and surface area, by difference between full geometry (yellow, which overlaps with blue along the cell body) and a geometry where the FPs are filtered out (blue). **(B)** Model used to identify the cell body of the cells, with expression for boundary condition. Once the system reaches steady state, regions within 90% of the maximum value are assigned as the cell body. **(C)** Measurements of lengths for major processes and branches. **(D)** Representative geometry of 25% of a cell. Rat podocytes (RP) used in this figure are numbered RP1, RP8, RP9, RP11, and RP13, respectively.

We clustered quantitative geometric parameters for podocyte morphology into three distinct compartments based on their reaction-diffusion dynamics: cell body (CB), major processes (MP) that includes primary and secondary processes, and FPs denoted with appropriate subscripts, respectively, plus the nucleus (Nu). The volume, V, and surface area, A, when without subscripts, represent the respective parameters for the whole cell. The CB, MP and FP volumes were derived from the segmented images. Different filters that were based on reaction-diffusion dynamics, allowed reconstruction of a whole cell (yellow surface in Fig 1A) and a “foot process-free cell” (blue surface in Fig 1A). The difference between the corresponding volume and surface areas for the full vs. FP-free cell allowed for estimation of FPs geometrical properties. We found that FP structures increased the cell volume by 20%, whereas more than doubling the cell surface area.

Boundaries of the different spatial components of the cell were estimated by applying a stratification method that is analogous to a well-known heat transfer problem (21). The primary processes of a podocyte have a large surface-to-volume ratio in comparison to its cell body. Therefore, a podocyte with uniform volumetric synthesis and diffusion of a given protein within its entire volume and with transport of protein to the extracellular space (proportional to local intracellular concentration) will equilibrate to a much lower concentration at the primary processes relative to the cell body (Fig 1B, analog heat transfer problem of uniform heat generation with convective boundary condition). Our analysis revealed that the cell body volumes range from 30 to 50% of the total cell volume; FPs correspond to 20% of the total volume; and the remaining 50% to 30% of the volume corresponds to primary and secondary processes. The distances between the center of the cell body and branch points, or ends of the processes, were measured using the filament tool in Imaris, as exemplified in Fig 1C. Consistent with the volumetric analysis, there was increase variance in process lengths and number of branches (Supplementary Material, Fig S1).

Since the branching parameters such as process length, counts, and distances were widely variable, we utilized the analytical descriptors of segregated volumetric units to construct a representative podocyte geometry. This analytical geometry was constructed with the objective of recapitulating the cell properties that are relevant for reaction diffusion equations (length, surface area, and volume) rather than visual similarity. The application of symmetry allowed us to reduce the mesh sizes needed for numerical discretization and the computational cost. FPs emanate from major processes in a symmetric fashion, generally with equal number of FPs in each side of the process of origin. Therefore, in our analytical geometry, the major processes are generated as halves, using the sagittal plane as a reflexive boundary condition. The FPs emanate perpendicularly to such boundary. Since there was no signature branching pattern for these cells, we also imposed axial symmetry. Consequently, the analytically constructed geometry produces simulations of a full podocyte cell, but at ¼ the size.

The representative geometry was initially built by taking into account only the cell body and major processes. Once the surface-to-volume (see Table 1) and distances of 18 ± 6 μm average distance from centroid to branch point, and 39 ± 2 μm for the three furthest endpoints, satisfied the analysis described above, the FPs were added. The quarter of the cell body (cell body plus major processes) has a volume of 420 μm^3^ and surface area 695 μm^2^. After 233 FPs were added, the volume corresponds to 530 μm^3^ and surface area of 1683 μm^2^ (Fig 1D). Table 1 demonstrates that the volumetric properties of the constructed geometry correspond to a good representation of the analyzed experimentally imaged cells. The surface area of the constructed FPs is on the upper range of the analyzed values. This is likely a conservative estimate; the resolution of the acquired images is of the length scale of the FPs, and a loss of surface detail is expected. The non-spatial computational model described below only uses volumetric variables, and is not affected by this assumption.

### Actin cytoskeleton

Our objective was to build a minimal ordinary differential equations (ODE)-based dynamical model that describes the exchange of actin monomers between monomeric (G-actin), fibrous (F-actin) and bundled states. Therefore, for this study, we omitted details such as actin ATPase activity, number and lengths of actin branches, or number of filaments per bundle. Different molecules with similar functions are treated implicitly. For example, a generic “bundling coefficient” is used rather than different expressions corresponding to individual molecular contributors of actin-associated proteins or crosslinkers for bundling. However, different parameters in the model can be indirectly related to the activity of Rac1 and RhoA, allowing us to use these as surrogates for the corresponding signaling pathways. Furthermore, the levels of actin polymer in the FPs are surrogates for their structural stability. Next, we show that analysis of this model provides novel insights on the how the balance of small GTPase activities affects the spatial stability of podocyte actin cytoskeleton.

### Non-spatial model reveals relationship between regulators and presence of stable points

Fig 2A shows a schematic representing the relationship between each state and the nomenclature used for parameters. In our model, we focused on the three discrete states of the actin cytoskeleton, namely monomeric G-actin, filamentous F-actin, and stress fibers or bundles, represented by Ga, Fa, and Bu, respectively. Monomers can dimerize to form *de novo* filaments (assuming nucleation factors are available) or add to fibers or bundles. The simplest mathematical representation is to have such rates directly proportional to the concentration of molecules in monomeric and fiber states, with macroscopic rate constants represented by the parameters Y_f_ and Y_b_. Dimerization (initiating new filaments) was assumed to be subject to the same rate constant as elongation, in order to reduce the total number of parameters and facilitate the analysis. This assumption does not impact the model analysis, as presented in greater detail in the supplementary material (Figs. S2-S3). The filament state is further enhanced by positive feedback (α_f_). This represents the molecular machinery that promotes polymerization in the FPs. It is a surrogate for the polymerization triggered by phosphorylation of nephrin and focal adhesion signaling through Nck or Rac1 for example (22, 23), and represented by a Hill function. Therefore the parameter α_f_ is non-zero only within the volume representing FPs. A non-linear functional form is necessary to represent both nephrin and small GTPase Rac1 driven polymerization (24-26). The denominator identifies the high sensitivity region (the positive feedback term is only significant if Fa is at least of the same order of magnitude as the parameter k), and ensures that the apparent rate constant is bounded. Activation of RhoA and cross-linking factors result in the formation of actin bundles. New bundles are formed by merging fibers, while existing bundles grow by addition of fibers (α_b_). Once a fiber becomes bundled, more crosslinks are added and it is unlikely to simultaneously break all connectors between a single fiber and the bundle it belongs to. However, there is turnover of monomers either from depolymerization or stress related rupture, both for fibers and bundles (β_f_, β_b_) (27, 28). Based on these considerations, equations 1-equations 3 below describe the exchange of actin monomers among the three states. We assume that the species Fa and Bu occur only within the FPs, while Ga freely diffuses to the different regions of the cell.

**Fig 2.**
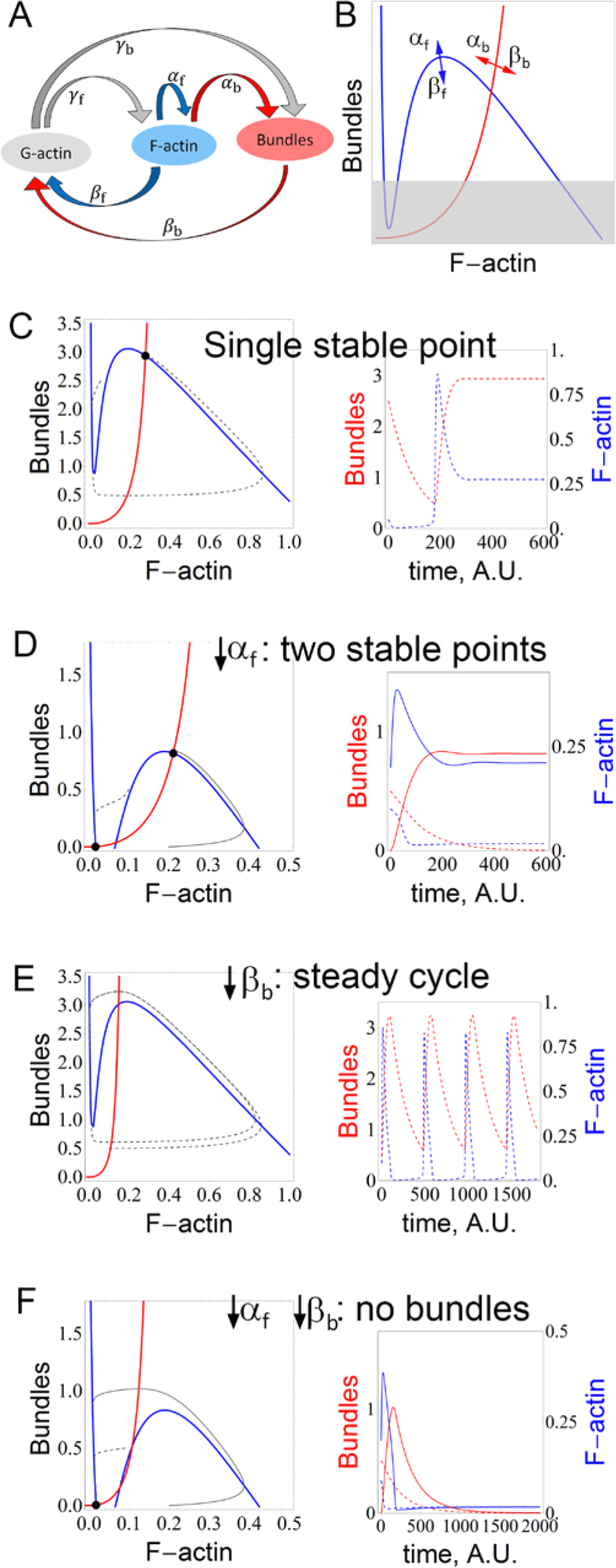
Minimal model for actin cytoskeleton in FPs. **(A)** Representative diagram and nomenclature for parameters. **(B)** Summary for relationship between nullclines and parameters α_f_, β_f_, α_b_ and β_b_. Blue curves are nullclines for Eq. 1 and red curves for Eq. 2. The gray shaded region represents the “effacement region” where FPs are unable to resist mechanical stress and lose their integrity. **(C)** A system with strong α_f_ (or weak β_f_) has a single stable equilibrium point. The concentrations for bundles and F-actin in the dashed trajectory of the phase plane (left, dashed gray line) are plotted in the time-series to the right, in red and blue, respectively. **(D)** Weak α_f_ or strong β_f_ give rise to a second stable equilibrium point, representing the collapse of bundles. The concentrations for the solid trajectory in the phase plane (left) are plotted with the solid lines in the time series (right). **(E)** Weak bundle turnover rate β_b_ or strong bundling α_b_ destabilizes the system, and there are no longer stable equilibrium points. However, the cyclic behavior might be able to keep the bundles sufficiently strong at all times. **(F)** A combination of weak bundle turnover rate β_b_ (or strong bundling α_b_) and weak positive feedback (α_f_) results in complete collapse of the actin cytoskeleton. Model parameters listed in Table S1.

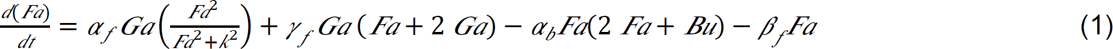

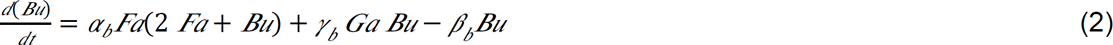

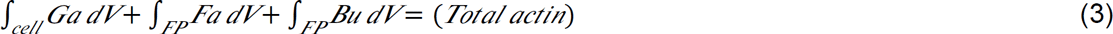

Since the exact concentrations of actin at each state in the podocyte are unknown, all of the concentrations in the above equations are set as non-dimensional, which would allow quantitative comparison of state spaces. It is known that the ratio of total amount of monomeric actin to polymerized actin in the healthy podocyte is about 1:2 (29).

The intersections between the nullclines (lines identifying values of variables that result in zero time derivative) for the variables F-actin and Bundles reveal the equilibrium points for the system, and increasing any of the parameters presented in Fig 2B will move each nullcline as indicated by the arrows. Bundles are able to sustain mechanical stress better than a filamentous network, and a minimum bundle density is expected to be necessary for each individual FP to persist (30). If the polymerization or bundling rates are not sufficient to overcome the turnover rates, the equilibrium point moves to the “effacement region” of the diagram that is demarcated with gray shading, where loss of FP integrity and specialized morphology (i.e., effacement) is expected.

### The behavior of the non-spatial model reconciles the varied experimental findings on the roles of Rac1 and RhoA

As illustrated by the two trajectories in Fig 2C, under conditions of a parameter regime with a single stable point, the FPs are able to sustain strong bundles irrespective of the initial conditions, or perturbation. This diagram represents ideal circumstances for a healthy stable podocyte. In contrast, a weakened α_f_ (due to decreased Rac1 activity, or defective Nck signaling) creates a second stable equilibrium point (Fig 2D). If the system moves to this new state, with negligible bundles, the FP integrity will most likely be lost (since this point is located in the effacement region of Fig 2B). Another unsatisfactory parametric set corresponds to increased bundling (α_b_), or even decrease in the turnover of the bundles (β_b_), destabilizing the system. Depending whether α_f_ is high or low, it may be subject to a cyclic behavior or collapse, respectively (Fig 2E-F). The results in Fig 2 correspond to a non-spatial model of actin dynamics, with FPs comprising 20% of the total volume (Eqs. 1-3).

The behaviors described by the model are consistent with the experimental observations for response to different activity levels of RhoA *in vivo.* A basal level of RhoA in podocytes (known to activate myosin and promote bundling) has been shown to be necessary for healthy glomerular function (19). Weak bundling (α_b_) would move the equilibrium point towards higher concentration of F-actin and lower concentration of bundles, shifting the equilibrium point towards the effacement region shown in Fig 2B. In this non-spatial model, the reasons for the damage caused by hyperactive RhoA is not obvious. Our model suggests that RhoA hyperactivity may lead to an imbalance between bundling and depolymerization. In this scenario, the bundles “consume” all the actin, and the minimum density of fibers required to feed the positive feedback is no longer achieved. Eventually the fiber density becomes too low and the bundle turnover surpasses its formation. Once this happens, the cytoskeleton may either be subject to terminal (Fig 2F) or temporary collapse (Fig 2E). As the bundles collapse, more monomeric actin becomes available. If the positive feedback for fiber is sufficiently strong, the fiber density recovers and the bundles density increases, resulting in cycles of weaker and stronger bundle density. The instances of weaker bundles may be sufficient to make the FP more susceptible to effacement under increased stress or even under physiological conditions. This scenario is further explored by spatial modeling in the next section.

A weaker positive feedback (α_f_) is the mathematical representation of a system where Rac1 is inhibited. The low α_f_-system in Fig 2D has a stable equilibrium point with moderate bundles. This result is a feasible mechanistic explanation for the observation that in healthy podocytes Rac1 may be inhibited. The presence of another stable point, which represents the collapsed cytoskeleton, demonstrates that this region of parameter space is not as robust as the one in Fig 2C. In addition, sustained high blood pressure (corresponding to increased β_b_ in the model), moves the bundle nullcline as indicated in Fig 2B, further decreasing the bundle concentration of the stronger stable point. Our model agrees with the physiological observation that, under such conditions, damage is observed (16).

### Diffusion limited process breaks the oscillatory synchrony, resulting in progressive localized damage

When our model is converted to a partial differential equations (PDE)-based dynamical model using the representative 3D podocyte geometry shown in Fig 1D, spatial heterogeneities in the oscillatory behavior emerge (Fig 3). This gives rise to permanent localized loss of bundles, which can be considered as a surrogate for effacement of FPs (Fig 3A-D). As monomeric actin is consumed, it must diffuse from its largest pool in the cell body to all FPs. The varying distance from this major G-actin source to the arrays of FPs, which act as individual G-actin sinks, produce gradients in G-actin, triggering asynchronies. Thus, bundles are collapsing in some FPs, while being strengthened in others. Consequently, as some FPs “release” their pool of G-actin, those monomers are sequestered by neighboring FPs, which are reciprocally reinforced. This is demonstrated at the ODE level in Fig 3E and Fig 3F which have identical kinetic parameters, but different total amounts of actin (70% and 115% of Fig 2E). Fig 3A-C show the bundle concentration at different elapsed time after α_b_ is increased. After elimination of some FPs, the remaining ones are able to build stronger bundles, temporarily enhancing their ability to overcome stress. However, the system is still unstable and damage slowly progresses. Examination of Fig 3C, in particular, shows that although this is a purely deterministic system, the spatial pattern of bundle loss among the FPs appears to be random. Furthermore, the precise value of the perturbation in α_b_ will produce different patterns. For example, different α_b_ will result in a different steady cycle amplitude range and frequency in the ODE model, which will be translated into a different pattern for the loss of synchrony in the spatial model. Such unpredictable behavior following a change in initial conditions is the hallmark of a chaotic system. Fig 3D shows how the asynchrony can be followed by permanent damage in some FPs. In some of the FPs, bundle concentrations drop permanently to 0, which would correspond to FP effacement. This example illustrates the potential impact of hyperactive RhoA on the podocyte integrity.

**Fig 3.**
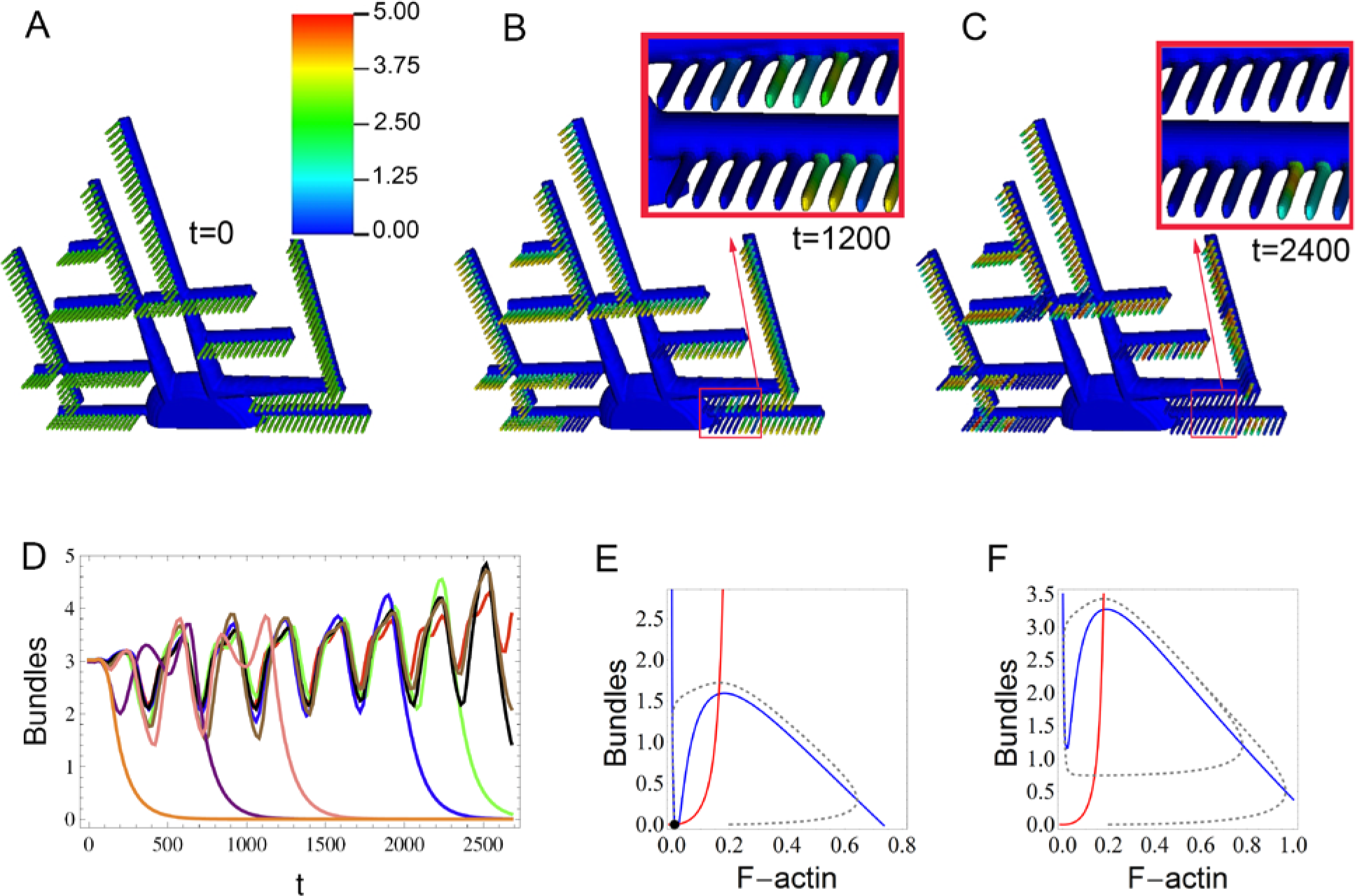
Impact of hyperactive bundling, α_b_, in FP integrity. Spatially, the cyclic behavior (triggered by sudden increase in α_b_ at t = 40) gives rise to loss of synchrony and progressive FP effacement (A-D). **(A)** Stead state bundle concentration for parameters as in Fig 2C. **(B)** Snapshots of bundle concentration in response to increased bundling, at time t = 1200 and **(C)** time t = 2400. Insets correspond to amplified and slightly rotated view of respective boxes. The colorbar represents bundle concentration. **(D)** Timecourse of bundle concentration at randomly picked FPs. ODE solution for increase in α_b_ predicts cytoskeleton collapse **(E)** when the actin pool is reduced (70% than in Fig 2) or stronger yet still unstable bundles **(F)** for systems with larger pools of actin (115%). In spatial simulations the positive feedback, α_f_, is localized to FPs only, and zero elsewhere.

### Localized or transient Rac1 hyperactivity can damage the podocytes

Each podocyte is subjected to a spatially diverse stimulus: it adheres to a segment of the coiled glomerular capillary vessel, and it interacts with several other podocytes. The cell-cell and cell-basement membrane interactions may lead to enhanced Rac1 activity, downstream of transmembrane slit diaphragm protein nephrin or focal adhesions (23, 31). Using our spatial model, we can demonstrate how this could lead to local damage. The ODE system with parameters as in Fig 2C has a single stable solution. However, this model is only valid if all parameters are uniform among all FPs. To explore and analyze the effect of differential actin bundling activities we enhanced the ODE model to include two compartments (Eqs. S2-S6). The monomer Ga is assumed to diffuse infinitely fast, resulting in the same concentration in the two compartments. The fraction “FP_1_” of the FPs in compartment 1, retain the original value of α_f_, while there is a localized increase in the positive feedback strength for the complementary fraction of FPs, “FP2”, in compartment 2. Fig 4A shows the steady state response for bundle concentrations in such an asymmetric cell, with constant positive feedback values α_f_ and α_f_+Δα_f_ for the fractions of FPs labeled FP_1_ (blue mesh) and FP_2_ (red mesh), respectively. As expected, for small values of Δα_f_, each fraction of FPs have a new concentration of bundles, weaker (FP_1_) or stronger (FP_2_) than when Δα_f_ is 0 (green line). As Δα_f_ and FP_2_ fraction increases, the bundles in the region FP_1_ collapse. Also note that as FP_2_ fraction increases, the total amount of actin becomes a limiting factor, and the local strength for the bundles cannot reach the high values observed in small FP_2_ fractions. This plot also illustrates that even though the machinery for F-actin polymerization is ubiquitous, the regions of the cell that are able to gather stronger polymerizing factors outcompete the remainder of the cell for the primary resource (G-actin), resulting in polarization of F-actin and bundles. This result is consistent with the observation that in podocytes, the actin cytoskeleton is present only on the FPs (32), where a high concentration of proteins that mediate actin polymerization are specifically localized (33).

**Fig 4.**
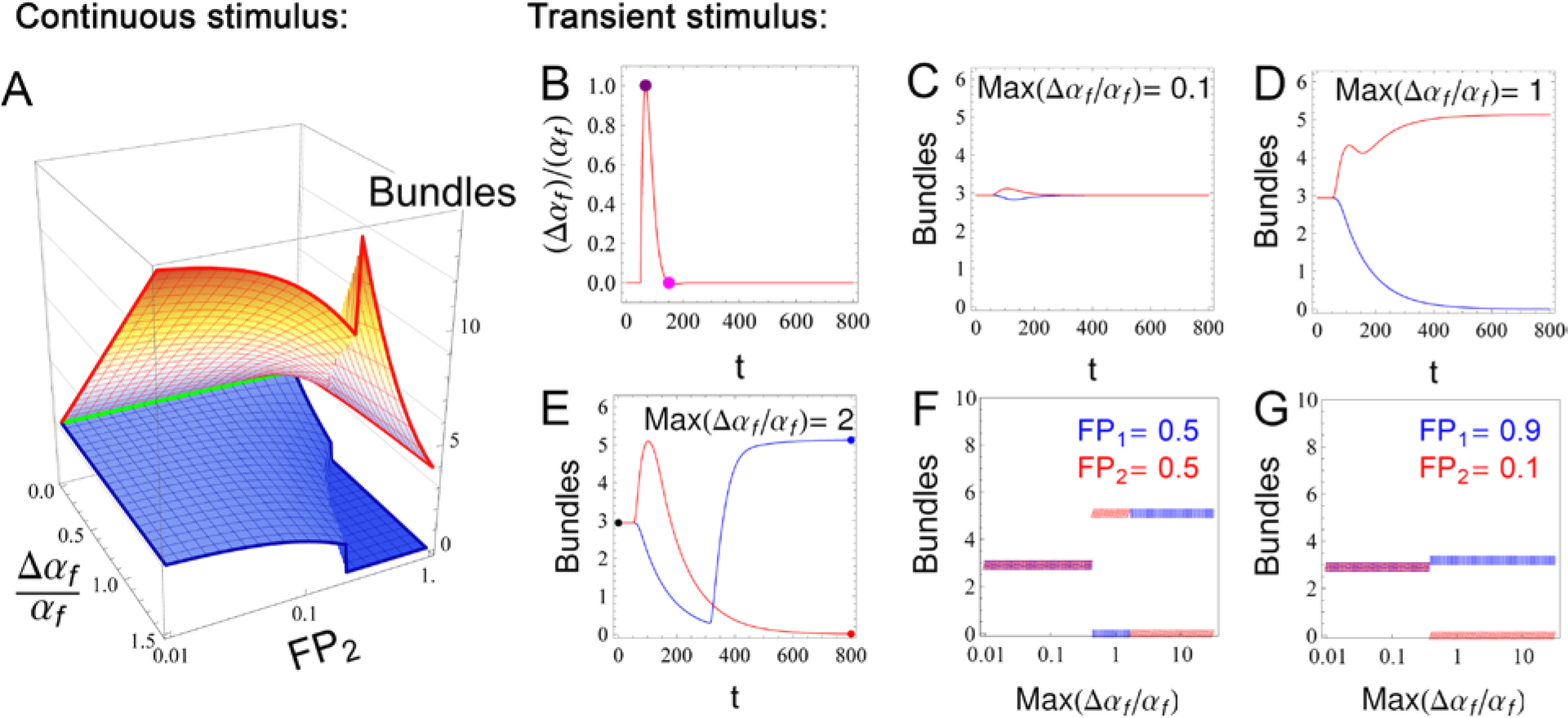
Impact of spatial inhomogeneity for positive feedback α_f_ in FP integrity. A fraction of the FPs (FP_2_) are subject to a stimulus that locally enhances actin polymerization by Δα_f_, whereas the remaining FPs (FP_1_) are under regular polymerization conditions, α_f_. This stimulus may either be sustained (shown in A) or transient (shown in B-G). **(A)** The fraction FP_2_ of the FPs has the positive feedback stronger by Δα_f_ than the fraction FP_1_ of FPs. When this difference is sustained, the bundle intensity (z-axis) in the fractions FP_1_ (blue mesh) and FP_2_ (red mesh) depend on the intensity Δα_f_ and the ratio of FP_2_ to FP_1_ (FP_1_ + FP_2_ = 1). The green line represents the intersection (Δα_f_ =0) for bundle concentration for regions FP_2_ and FP_1_ for the corresponding values of Δα_f_ and FP_2_ (notice that the axis for FP_2_ is in log scale). As Δα_f_ increases, FPs with stronger feedback form stronger bundles. If the fraction of FPs with enhanced feedback (FP_2_) is small, the FPs with normal α_f_ (FP_1_) are unperturbed, while the bundles in FP_2_ are strengthened. Because there is a limited total amount of actin, in order to get stronger bundles in FP_2_, the bundles in FP_1_ must loose actin. As the intensity Δα_f_ and region FP_2_ of enhanced feedback get larger, the weakening or collapse of FPs with original feedback strength (FP_1_) is observed. In A the value of Δα_f_ is constant in time. In C-E the value of Δα_f_ follows the timecourse as in B, with varying intensities as indicated in C-E. Equal volume fractions for FP_1_ and FP_2_ were used, with red curves corresponding to FP_2_ and blue to FP_1_. Depending on the intensity of the transient stimulus, the region FP_2_ may recover the same bundle concentration as in FP_1_ (C), develop permanently stronger bundles (at the expense of FP_1_, D) or collapse (E). F. The steady state values for concentration of bundles in fractions FP_1_ (blue) and FP2 (red) are shown as a function of stimulus intensity, consistently with C-E. Different fractions of FP_1_ and FP_2_ will impact the steady state values (G and Fig S5).

Fig 4C-E show the impact of a localized transient enhancement (shown in Fig 4B) of the positive feedback on region FP_2_ only (50% of the FPs). If the stimulus is weak, both FP_1_ and FP_2_ regions recover the uniform bundle strength (blue and red lines, respectively, in Fig 4C). For moderate stimulus, FP_1_ fraction loses bundles permanently and FP_2_ reaches a new steady state, with stronger bundles (Fig 4D). For very strong Δα_f_, the initial response is as expected, but at longer times, the bundles in FP_2_ collapse while the bundles in FP_1_ are enhanced Fig 4E). The phase-plane diagrams in Fig S4 help explain the switch. At time zero, either FP_1_ or FP_2_ fraction has the same composition regarding bundles and F-actin. With strong Δα_f_, FP_2_ fraction consumes G-actin in order to develop more and more fibers, while the production rate of fibers for FP_1_ becomes weaker than its turnover. At this point, the two regions of FPs are asynchronous, and their volume fractions and proportions of F-actin and bundles (i.e., their position in the phase-plane) will determine which fraction “wins over” the available G-actin (note the trajectories in Fig 2C). In summary, the steady state response to a localized transient peak in the polymerization positive feedback can either lead to both regions of the FPs to recover the original uniform bundle concentration, or lead to one of the stimulated (FP_2_) or unstimulated (FP_1_) regions to collapse (Fig S5).

The same range of responses was observed in our spatial simulations. Consistent with the results in Fig 4C-G, the spatial simulations revealed that localized transient increase in the positive feedback α_f_ may lead to localized damage of FPs both in the regions within and adjacent to the stimulus (Fig 5). Time zero has all FPs in steady state and at same bundle concentration (as in Fig 3A). After time 40, only the region FP_2_ (as marked in Fig 5) is subject to a transient increase in the parameter for positive feedback α_f_, while all other FPs are subject to constant α_f_ (region FP_1_). The timeseries plot in Fig 5A indicates the bundle concentrations in FPs identified by the corresponding colored arrowheads. Because the availability of monomeric actin is diffusion limited, a gradient of responses is observed. Within the region FP_2_, there are FPs that recover the original bundle concentration once the stimulus is removed (e.g., the FP indicated by the orange arrow head in Fig 5A). There are also FPs that collapse within the same region (red arrowhead). Interestingly, it is the FPs near the boundary that permanently lose their bundles. Similarly, among the FPs in close proximity to FP_2_ (yet with constant α_f_), some also permanently collapse (blue arrowhead). The 3-D movie of Fig 5 is in Supplementary Movie 2. In the movie, the bounding box identifies the region FP_2_. A second example is shown in Supplementary Movie 3, with a significantly larger region for FP_2_, comprising all FPs within the bounding box.

**Fig 5.**
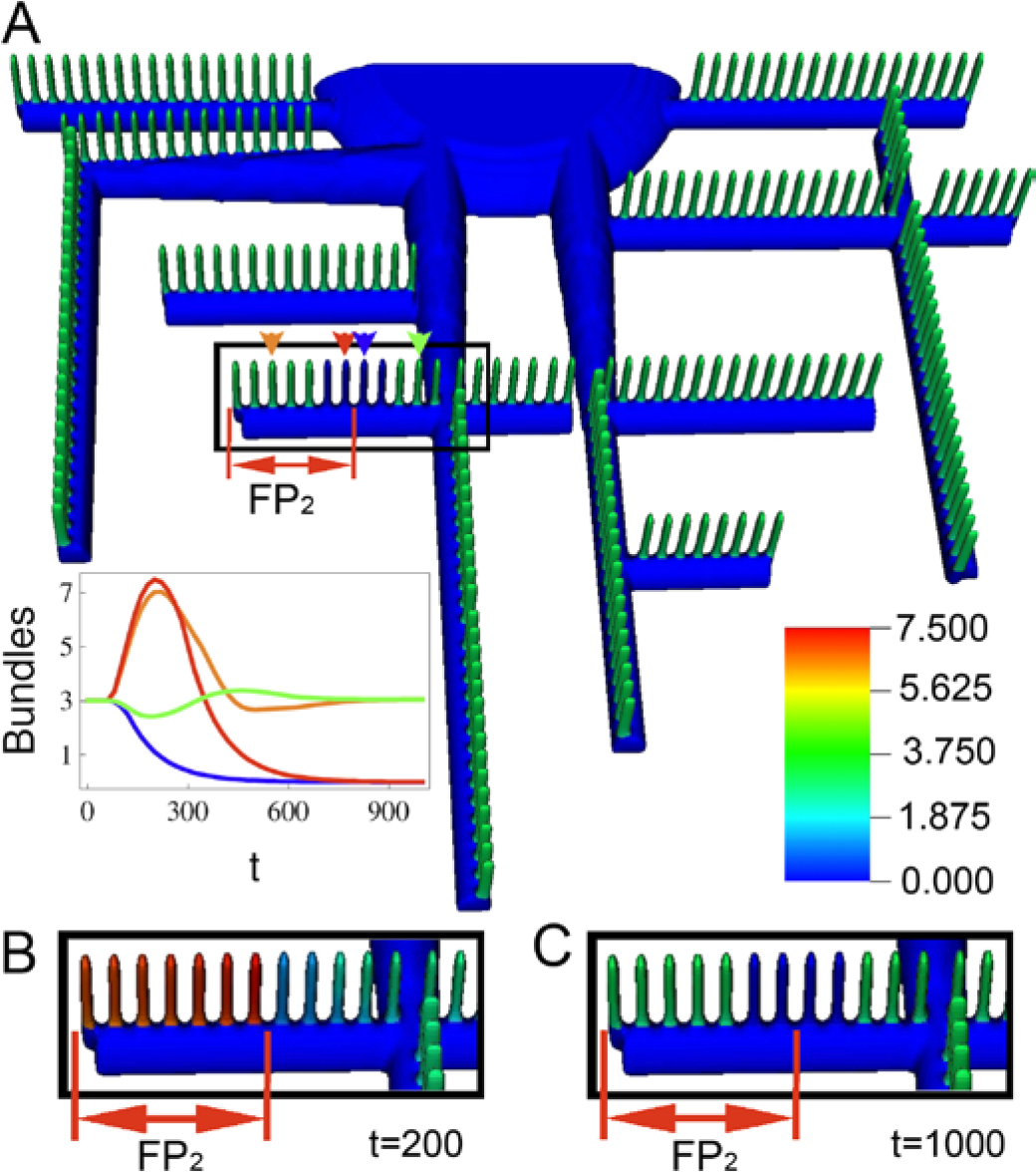
Spatial simulation showing impact of transient localized increased positive feedback α_f_ in FP integrity. **(A)** State state bundle concentrations after transient stimulus in α_f_ in region FP_2_. The transient responses for the specific FPs identified by arrowheads are plotted using the corresponding colors in the inset. Zoomed snapshot of the highlighted region is shown at **(B)** time = 200, and **(C)** time = 1000.

### Compensatory mechanisms: the earlier the better

Next, we studied the impact of decreasing bundle turnover rate β_b_ and the potential ways to regulate podocyte FP integrity. Bundle turnover rate, β_b_, accounts for depolymerization and damage of bundles due to mechanical stress in the glomerulus. It is expected that under high or low blood pressure, β_b_ would be enhanced or attenuated, respectively. As described above, cytoskeletal stability is dictated by the relationship between the F-actin and bundles nullclines, and as illustrated in Fig 2B, different parameters move the stable equilibrium points in different directions. Initially, the healthy podocyte is in steady state in its single equilibrium point (Fig 6A). By decreasing the bundle turnover rate, the system is destabilized and progressive damage and effacement of FPs are observed (Fig 6B-C). Here in this complex geometry, the interplay of eq.1 and eq. 2, through diffusion of G-actin, produce marked spatial heterogeneity in the pattern of bundles. The trivial compensatory mechanism would be to restore the original value of β_b_ after a certain amount of time Δt_1_ (Fig 6D). Most of the FPs that survived up to the time of the correction (Fig 6B) would have reached stability after elapsed time Δt_2_ (Fig 6E). A similar result is observed if the compensatory mechanism is applied at a later time (Fig S6). The permanent effacement of several FPs provides a larger pool of actin to be incorporated by the surviving ones, resulting in stronger bundles. This would correspond to the loss of some FPs and the strengthening of others. The timecourse for the bundles in the FPs highlighted by solid arrows is plotted with corresponding solid lines in Fig 6H. The gray arrowhead identifies the time point used for the 3-D snapshots. These spatially asynchronous and irregular timecourses further substantiate the chaotic tendencies of this system.

A second alternative for a compensatory mechanism is the decrease of α_b_, representing decreased bundling coefficient or concentration of cross-links (Fig 6F). We show that this mechanism can potentially achieve equivalent success as the trivial case of restoring β_b_. Once again, the sooner the intervention, the smaller the population of damaged FPs (Fig 6F-G, and timecourse of FPs identified by dashed arrows represented in dashed lines in Fig 6H and Fig S6).

**Fig 6.**
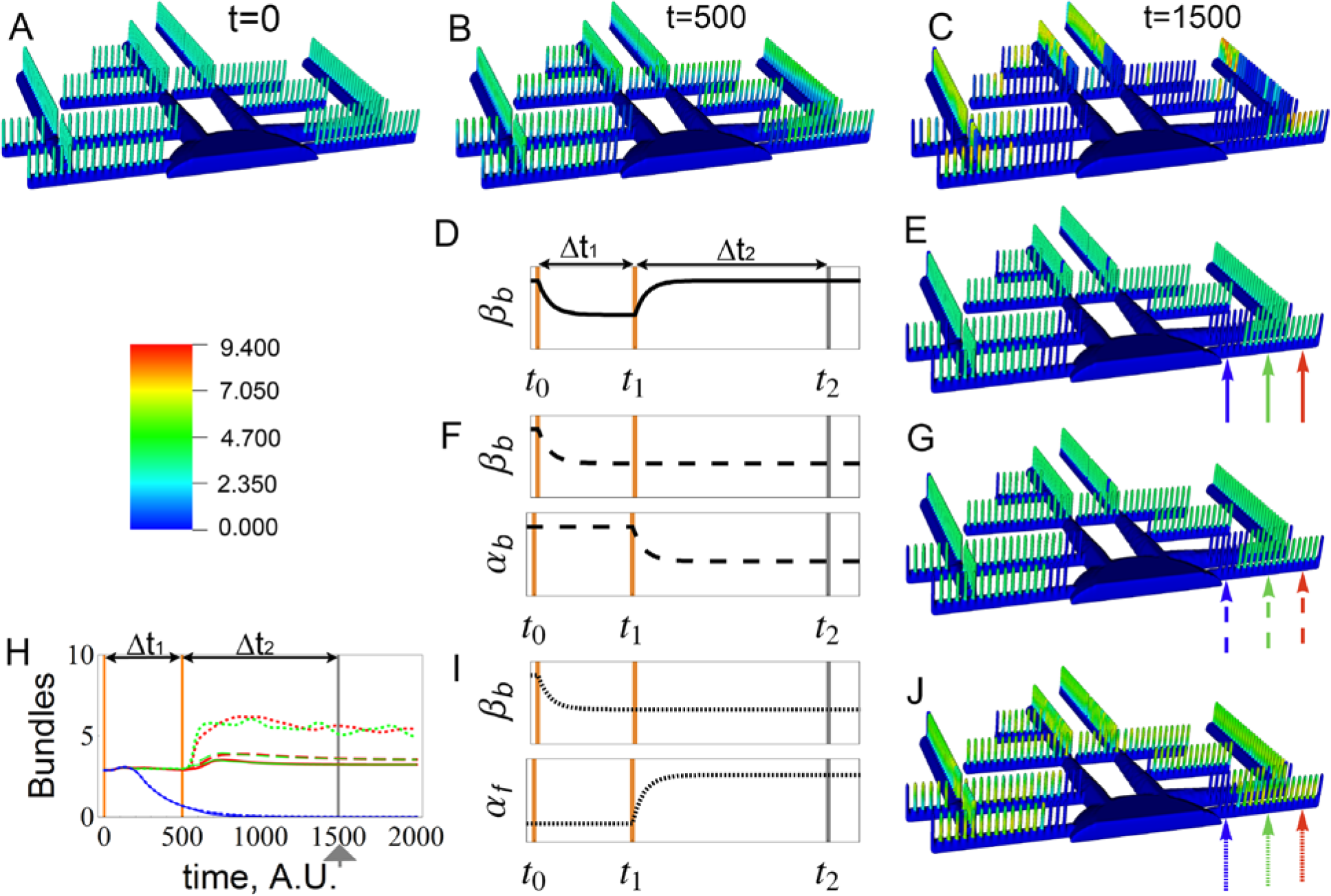
Compensatory mechanisms for podocyte disruption. A. Bundle density for unperturbed podocyte. Progressive loss of FPs due to decrease of β_b_ imposed at time t = 0, snapshots at time **(B)** t = 500, and **(C)** t = 1500. We predict that FP effacement can be halted by three different mechanisms that is systematically introduced at time t_1_. **(D)** Recovery of β_b_ at time t_1_ results in **(E)** stabilization of great majority of the remaining foot process. **(F)** Decrease of α_b_ at time t_1_ while holding β_b_ constant results in **(G)** similar stabilization. Finally, **(I)** increase of α_f_ while holding β_b_ constant **(J)** produces similar spatial results. All three interventions prevent progressive effacement (compare C with E, G and J). **(H)** Timecourses for spatial average of bundle concentration in the FPs identified by arrows in snapshots E, G and J (at time 1500, gray arrowhead). Linestyle follows the same pattern as arrows. The same color scale is used for all the 3-D snapshots.

Finally, we also explored the impact of increasing the positive feedback, α_f_, corresponding to hyperactivating Rac1 or other polymerization signals coming from the slit diaphragm (Fig 6I-J). Now high level of polymerized actin is maintained in each of the surviving FPs, however there is a new mode of oscillation for bundles (and F-actin) along each FP. The simulations suggest that if applied early enough, this compensatory mechanism may help keep the podocyte attached (Fig 6J). Similarly to the previous cases, the surviving FPs are able to build stronger bundles, and in spite of the oscillatory behavior, equivalent number of FPs seem to be preserved in comparison to the previous examples (Fig 6J, Fig 6E, Fig 6G). The overall conclusion is that there are several potential compensatory regulatory mechanisms that may be activated to restore FP stability, and the sooner the regulatory response is activated, the larger the number of surviving FPs.

## Discussion

We acquired 3-D electron microscope imaging data of rat kidney glomeruli, individually segmented podocytes, and quantitatively analyzed their morphological features to determine the average volume and surface area of the cell body, major processes, and foot processes. This resulted in construction of a representative 3-D podocyte model that is amenable to PDE-based reaction-diffusion modeling with physiologically relevant mechanisms. To the best of our knowledge, this is the first time such a detailed quantitative reconstruction was ever achieved. Through our morphometric analyses, we show that the FPs corresponded to 20% of the volume of the podocyte, while doubling its surface area. Consequently there is a large pool of diffusive monomeric G-actin that is available for the FPs. However, the G-actin pool is not instantly or uniformly available: the monomers must diffuse across the different regions of the cell. Our representative podocyte geometry carefully preserved the length-scales, volumetric and diffusive properties, quantity and patterns of FPs, primary processes, and cell body, thus enabling physiologically relevant simulations.

To explore the consequences of the unique podocyte morphology on the maintenance of cytoskeletal integrity of the FPs, we developed a minimal model of actin dynamics and compared non-spatial (ODE-based) simulation results with full reaction-diffusion (PDE-based) models in the constructed geometry. The minimal model represents key regulatory mechanisms controlling actin polymerization and fiber bundling. It contains positive feedback in actin polymerizations, representing Arp2/3-dependent branching nucleation on preformed mother filaments. This positive feedback results in regions of parameter space that produce oscillatory F-actin and bundle kinetics in the ODE simulations (Fig 2). Another mechanism that would be similarly represented by this positive feedback is actin polymerization downstream of nephrin phosphorylation and Nck localization to the FPs slit diaphragm.

Spatial simulations explicitly consider the additional process of diffusion of G-actin in and out of FPs from the large reservoir of the cell body and primary processes. The interplay of this diffusion and the positive feedback inherent in the actin dynamics can produce sharp regional differences in the level of actin bundling within closely neighboring FPs. Without bundled actin, the FP would be resorbed into the parent process leading to effacement. Therefore, we consider the loss of bundles in FPs as a surrogate for the loss of structural integrity. Even when a change in bundling activity is distributed uniformly throughout the cell, an apparently chaotic, spatiotemporal pattern of bundle formation and collapse is observed in the FPs (Fig 3). One of the major unanswered questions in podocyte biology is the source of the inherent heterogeneity in podocytopathies. Diseases, such as focal segmental glomerular sclerosis, affect only a subpopulation of podocytes with a large spatial variability in disease etiology. Based on our spatial dynamical model simulations, we suggest that even global changes in stress (e.g., due to hypertension) can lead to selective effacement of FPs with large spatial variability, which may explain the pathophysiological disease progression of numerous podocytopathies.

The balance of Rac1 and RhoA activity (that is represented respectively by actin polymerization and bundling in the model) needs to be tightly regulated to maintain FP stability. Stress (whether focal or global, transient or continuous) can lead to spatially heterogeneous chaotic behavior, and ultimately, to irregular heterogeneous patterns of bundle collapse in FPs. Fig 4 shows how transient activation of Rac1 (i.e., actin polymerization) in one collective region of FPs can shift the steady state balance between F-actin and bundles in all regions. A much more localized transient perturbation, as shown in the spatial simulation of Fig 5, can produce *permanent* changes in the bundle distribution within the system, both within the perturbed region and its immediate vicinity.

Direct damage to FPs, due to low blood pressure for example, can be captured in the bundling turnover parameter β_b_. Changing β_b_ globally, but transiently, can produce permanent changes in bundle distribution (Fig 6A-C); the severity of FP effacement (represented by loss of bundles) is directly related to the duration of the perturbation. It is possible that regulatory mechanisms could be modulated to alter Rac1 and RhoA activities in response to stress to ameliorate the imbalance. The behavior of the system under three potential compensatory interventions in response to lowered β_b_ are explored in Fig 6D-J.

The model for the actin cytoskeleton of the podocyte FP provides a framework for understanding recent findings on the *in vivo* activity levels of the small GTPases RhoA and Rac1. Our model captures the need for a balance between polymerizing, bundling, and turnover rates for the actin cytoskeleton. Our results are consistent with the observations that a minimum level of RhoA is necessary, however, hyperactivity results in destabilization and progressive loss of FPs. We propose that upon inhibition of Rac1, the cell may sustain its FP integrity with sufficient levels of actin bundles; however, the weakened positive feedback for polymerization gives rise to a second equilibrium point for the system, losing its robustness. It is also important to highlight that nephrin stimulated actin polymerization could also influence the parameter α_f_, and mutations on crosslinks would impact the parameter α_b_.

We hypothesize that healthy cells present a set of parameters that result in a single equilibrium point, representing FPs with strong bundles. In such circumstances, all FPs are stable and have identical properties. Using a reconstructed geometry, we studied the impact of parametric inhomogeneity. As demonstrated here, G-actin availability was a limiting factor and strong localized polymerization may disrupt the cytoskeleton of FPs elsewhere. The diffusion limited G-actin availability may also disrupt otherwise synchronized oscillations. This results in slow and progressive loss of bundles, the surrogate for effacement of FPs. Of course, the sudden localized collapse of actin structures may be driven by a local inhibitory effect (either decreased polymerization rate constant or enhanced turnover). However, in this work we demonstrate the feasibility of an alternative hypothesis: enhancing polymerization in remote regions of the podocyte, by a number of signaling pathways, may sufficiently disrupt the balance of G-actin availability to irreversibly drive the local collapse of existing FPs.

## Acknowledgements

This work was supported by funding from NIH (TR01 DK087650) to R Iyengar, JC Hone, and LM Loew, and from NIH (P41 GM103313) to LM Loew. EU Azeloglu is a NephCure Kidney International - ASN Foundation for Kidney Research scholar. We would like to acknowledge the Virtual Cell software team (Jim Schaff, Frank Morgan, Ed Boyce, Diana Resasco, Gerard Weatherby and Fei Gao) for their help with simulations; Ann Cowan for her help with Imaris image processing; Chiara Mariottini and Pedro Martinez for their help with 3-D reconstruction of podocytes.

